# EphrinB1 modulates glutamatergic inputs into POMC neurons and controls glucose homeostasis

**DOI:** 10.1101/2020.02.10.941765

**Authors:** Manon Gervais, Alexandre Picard, Bernard Thorens, Sophie Croizier

**Affiliations:** Center for Integrative Genomics, University of Lausanne, Lausanne, Switzerland

**Author notes:** **Correspondance**: Sophie Croizier, Ph.D, Center for Integrative Genomics, University of Lausanne, Genopode building, 1015 Lausanne, Switzerland.

**Keywords:** Hypothalamus, glutamatergic inputs, POMC neurons, ephrin, glucose homeostasis, feeding behavior

## Abstract

Proopiomelanocortin (POMC) neurons are major regulators of energy balance and glucose homeostasis. In addition to being regulated by hormones and nutrients, POMC neurons are controlled by glutamatergic input originating from multiple brain regions. However, the factors involved in the formation of glutamatergic inputs and how they contribute to bodily functions remain largely unknown. Here, we show that during the development of glutamatergic inputs, POMC neurons exhibit enriched expression of the *Efnb1* (EphrinB1) and *Efnb2* (EphrinB2) genes, which are known to control excitatory synapse formation. *In vitro* silencing and *in vivo* loss of *Efnb1* or *Efnb2* in POMC neurons decreases the amount of glutamatergic inputs into these neurons. We found that mice lacking *Efnb1* in POMC neurons display impaired glucose tolerance due to blunted vagus nerve activity and decreased insulin secretion. However, mice lacking *Efnb2* in POMC neurons showed no deregulation of insulin secretion and only mild alterations in feeding behavior and gluconeogenesis. Collectively, our data demonstrate the role of ephrins in controlling excitatory input amount into POMC neurons and show an isotype-specific role of ephrins on the regulation of glucose homeostasis and feeding.

## Introduction

Obesity and associated diseases, such as type 2 diabetes, are major public health concerns, and their worldwide prevalence are increasing at an alarming rate. Energy and glucose homeostasis are centrally controlled by complex neuronal networks that involve two main antagonistic neuronal populations in the arcuate nucleus of the hypothalamus (ARH): the anorexigenic proopiomelanocortin (POMC) neurons and the orexigenic Agouti-related peptide (AgRP)/neuropeptide Y (NPY) coexpressing neurons (1–4). In addition to the key role they perform in controlling feeding, POMC and AgRP/NPY neurons are involved in the control of glucose homeostasis (5, 6). Indeed, insulin action on AgRP neurons suppresses hepatic glucose production (7, 8) and chronic activation and inhibition of POMC neurons represses and stimulates gluconeogenesis, respectively (9). Moreover, alterations in POMC signaling, circuits, or neuronal survival are associated with disruption of glucose homeostasis. (10–12).

POMC and AgRP/NPY neurons are major integrators of peripheral hormones (insulin, leptin, and ghrelin) and nutrients (glucose) to control energy and glucose homeostasis through neuroendocrine and autonomic responses (13–17). Besides, POMC and AgRP/NPY neurons also receive abundant central information through GABAergic (inhibitory) and glutamatergic (excitatory) inputs (18). Notably, POMC neurons primarily receive glutamatergic inputs, whereas AgRP/NPY neurons receive primarily GABAergic inputs (19). However, the mechanisms underlying the development of POMC neuronal circuits and particularly the formation of glutamatergic inputs and how they contribute to glucose homeostasis and energy balance remain elusive.

During the development of the nervous system, cues that guide development orchestrate neuronal wiring to allow growing axons to reach their targets and establish synaptic contacts to build functional neuronal networks. EphrinB molecules form a family of cell-contacting proteins that specifically interact with EphA and EphB receptors to stabilize glutamatergic synapses, to recruit AMPA and NMDA receptors and to control the number of glutamatergic synapses. In the rat ARH, synapse formation begins postnatally and gradually increases until adulthood (20).

Here, we employed a transcriptomic approach to reveal that the *Efnb1* and *Efnb2* gene products (EphrinB1 and EphrinB2) are enriched in POMC neurons when glutamatergic inputs occur. *In vitro* approaches showed control of the number of excitatory inputs into POMC neurites by EphrinB1 and EphrinB2. In mice, the lack of *Efnb1* or *Efnb2* decreases the amount of glutamatergic inputs into POMC neurons but has isotype-specific metabolic effects impacting glucose tolerance, vagus nerve activation and insulin secretion (*Efnb1* deletion) or feeding and gluconeogenesis (*Efnb2* deletion). Our data show that POMC neurons belong to a complex neuronal network and can integrate central and peripheral information to control glucose homeostasis.

## Results

### Onset of glutamatergic inputs into POMC neurons

To visualize the development of excitatory presynaptic terminals in POMC neurons, we performed immunohistochemical labeling of presynaptic glutamatergic vesicular transporter (vGLUT2) in POMC neurons expressing a tdTomato reporter (*Pomc*-Cre;tdTomato) in postnatal days P6, P14 and P22 male mice (Figure 1A). At P6, glutamatergic inputs were already observed to be in contact with POMC neurons, and the amount of inputs increased gradually until P22 (Figures 1A-C).

**Figure 1.**
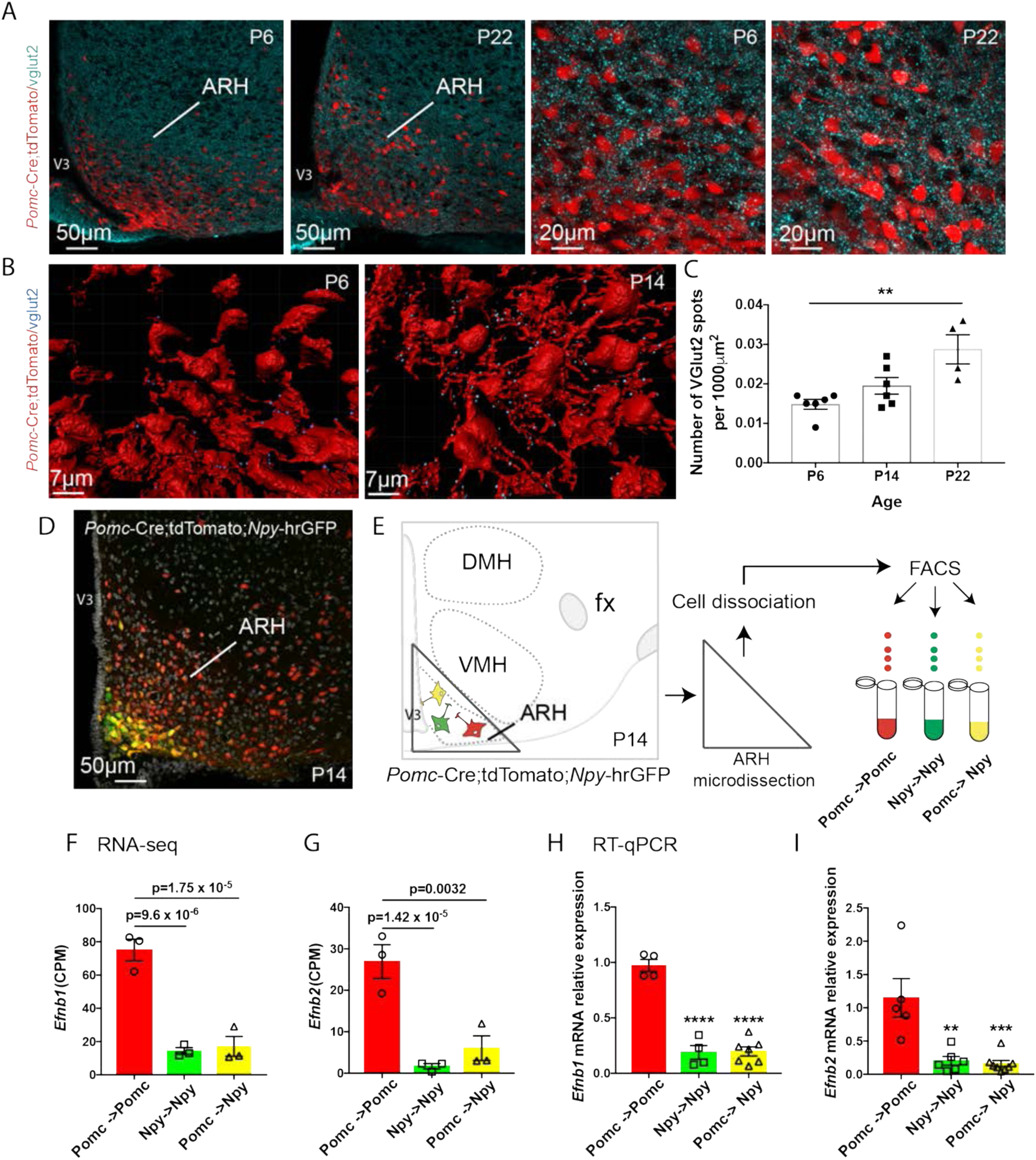
Enrichment of *Efnb1* and *Efnb2* mRNA in POMC neurons during postnatal development of glutamatergic inputs. (A) Confocal images and high magnification of vGLUT2-positive terminals (turquoise) into *Pomc*-Cre;tdTomato neurons (red) in the ARH of P6 and P22 male mice. (B) Example of 3D reconstruction (P6 and P14, IMARIS) and quantification (C) of vGLUT2-positive inputs in direct apposition with *Pomc*-Cre;tdTomato neurons of P6, P14, and P22 male pups (n=2-3/group, 2 sections/animal). (D, E) Image and schematic illustrating *Pomc*-Cre;tdTomato progenitors (Pomc->Pomc, red) and their partial co-expression with *Npy*-hrGFP neurons (Pomc->Npy, yellow) in the ARH of P14 male mice. *Npy*-hrGFP neurons not deriving from Pomc progenitors (Npy->Npy) are labeled in green. These three sub-populations are sorted by flow cytometry based on their fluorescence (E). RNA-seq data (Count per million, CPM) highlighted *Efnb1* (F) and *Efnb2* (G) genes as enriched in Pomc->Pomc neurons when compared to Npy->Npy and Pomc->Npy. *Efnb1* (H) and *Efnb2* (I) mRNA expression in Pomc->Pomc, Npy->Npy and Pomc->Npy populations measured by RT-qPCR. Data are shown ± SEM. Statistical significance was determined using One Way ANOVA test (C, H, I). \*\**P* ≤ 0.01 *versus* P6 (C), *versus* Pomc*-*>Pomc (I); \*\*\**P* ≤ 0.001 *versus* Pomc*-*>Pomc (I); \*\*\*\**P* ≤ 0.0001 versus Pomc*-*>Pomc (H). ARH, arcuate nucleus of the hypothalamus; DMH, dorsomedial nucleus of the hypothalamus; fx, fornix; VMH, ventromedial nucleus of the hypothalamus; V3, third ventricle.

Based on this developmental observation, we next performed transcript profiling of POMC-expressing (POMC^+^) and NPY^+^ cells at P14 to identify putative genes involved in glutamatergic synapse formation (Figures 1D and 1E). POMC progenitors can give rise to NPY neurons or POMC neurons. We thus performed RNA-sequencing (RNA-seq) experiments on POMC neurons (*i.e.,* POMC->POMC), on NPY neurons derived from POMC progenitors (*i.e.,* POMC->NPY), or on NPY neurons not derived from POMC progenitors (*i.e.,* NPY->NPY). These three subpopulations can be separated based on their fluorescence when using *Pomc*-Cre;tdTomato;*Npy*-hrGFP animals (red cells: POMC->POMC; yellow cells: POMC->NPY cells; and green cells: NPY->NPY) (Figures 1D-G, S1A and S1B). Principal component analysis (PCA) of the RNA-seq data did not distinguish POMC->NPY from NPY->NPY neuronal subpopulations (Figure S1A). We therefore only focused on results obtained from POMC->POMC and NPY->NPY neurons. This analysis revealed that 1586 genes were upregulated, and 1202 genes were downregulated in POMC->POMC neurons compared to NPY->NPY neurons. However, when the analysis was restricted to genes involved in axon guidance (Figure S1B, Table S1), out of the 37 identified genes, 2 were significantly upregulated and 4 were downregulated in POMC neurons compared to NPY neurons. In particular, *Efnb1* and *Efnb2* were 5.2- and 16-fold enriched, respectively (Figures 1F, 1G and Figure S1B). We further confirmed this increase in *Efnb1* and *Efnb2* mRNA expression in POMC neurons using RT-qPCR and observed consistent results; there was a 5.2- and 5.8-fold increase in *Efnb1* and *Efnb2* mRNA, respectively (Figures 1H and 1I). To determine whether *Efnb1* and *Efnb2* were differentially expressed in POMC neurons based on their anatomical location, we performed *in situ* hybridization and quantified the number of fluorescent punctate signals in *Pomc*-GFP^+^ neurons in distinct anteroposterior parts of the ARH of P14 animals (Figure S1C-H). This analysis revealed that POMC neurons homogeneously expressed *Efnb1* and *Efnb2* throughout the entire ARH (Figures S1E-H).

### *Efnb1* (EphrinB1) and *Efnb2* (EphrinB2) regulate the amount of glutamatergic inputs into POMC neurites

To study whether EphrinB1 and EphrinB2 can directly modulate the number of glutamatergic terminals on POMC neurites, we had to identify the glutamatergic area innervating the POMC neurons. Previous monosynaptic retrograde mapping showed that POMC neurons receive inputs from several sites, such as the preoptic area, the bed nucleus of the stria terminalis, the lateral septum and, in particular, from the PVH (Wang et al. 2015). The PVH is known to contain glutamatergic neurons (17); however, whether glutamatergic PVH neurons innervate ARH POMC neurons remain elusive.. To address this question, we used a retrograde viral approach using modified rabies virus SADΔG-mcherry combined with Cre-dependent helper adeno-associated virus (AAV) (Figure S2). An injection of these viruses into the ARH of *Pomc*-Cre male mice allows a retrograde monosynaptic spread from POMC neurons. We combined the detection of retrogradely labeled mCherry-positive cells with *in situ* hybridization of *vglut2* mRNA. This allowed us to visualize PVH cells that were retrogradely labeled with mCherry and were glutamatergic (Figure S2). These observations confirmed that POMC neurons receive glutamatergic inputs from the PVH.

We next examined whether ephrin receptors EphB1, EphB2, EphB3, EphB4, EphA4 and EphA5 were expressed in the PVH when glutamatergic terminals developed in POMC neurons (from P8 to P18). We found that *Ephb1*, *Ephb2* (Figure 2A), *Ephb3*, *Ephb4*, *Epha4* and *Epha5* mRNA (Figure S3) were expressed in the PVH during this postnatal period. *Ephb1* and *Ephb2* are well-known receptors of EphrinB1 and EphrinB2 (21). We thus assessed by *in situ* hybridization whether *Ephb1* and *Ephb2* mRNA were specifically expressed by glutamatergic neurons of the PVH at P14. Our results show that 97.2% and 88.2% of glutamatergic neurons in the PVH expressed *Ephb1* and *Ephb2*, respectively (Figures 2B and 2C). These data suggest that the establishment of PVH glutamatergic input into POMC neurons may depend on presynaptic EphB receptors and postsynaptic EphrinB (Figure 2D).

**Figure 2.**
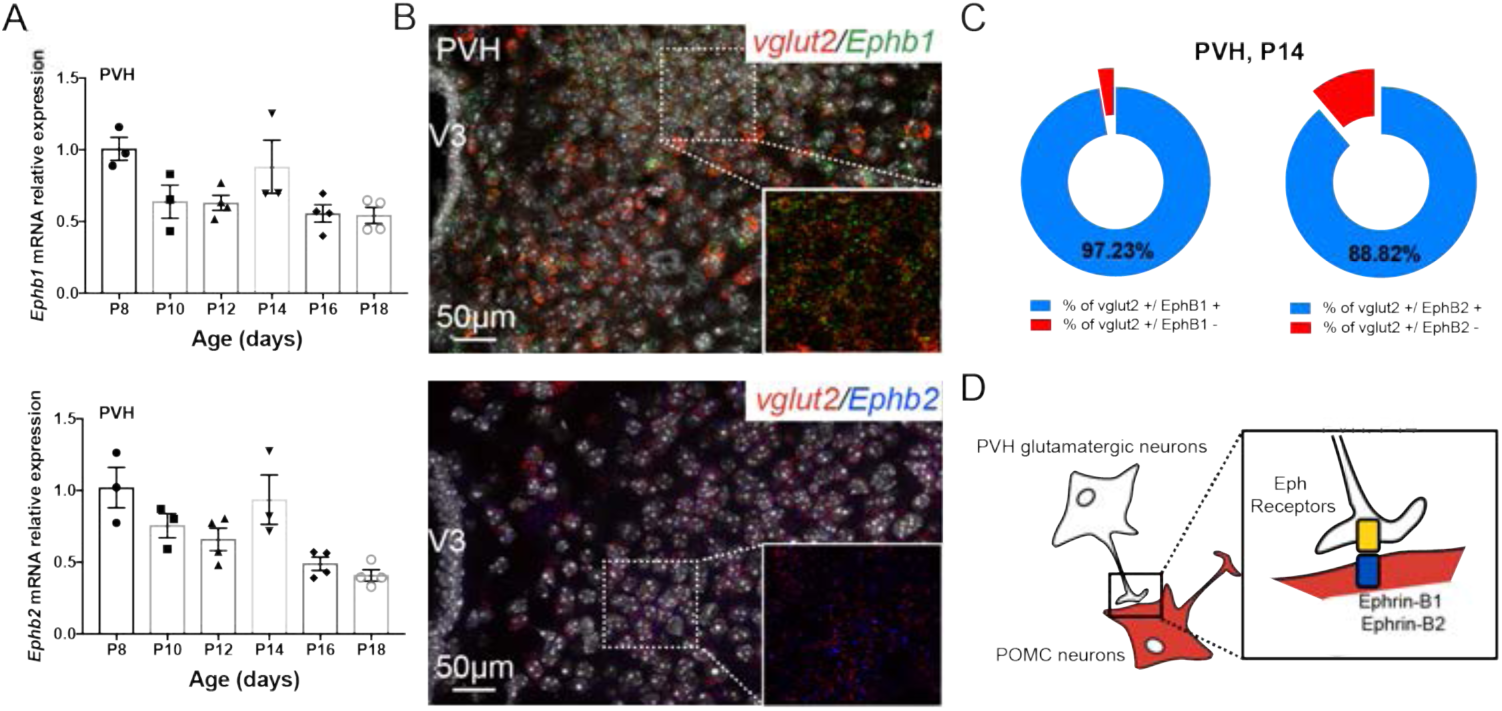
EphrinB signaling. (A) Quantification of *Ephb1* and *Ephb2* mRNA relative expression in the PVH of P8 to P18 male mice (n=3-4/group). Photomicrographs (B) and quantification (C) showing co-expression of *Ephb1* (green) and *Ephb2* (blue) mRNA with *vglut2* mRNA (red, glutamatergic marker) in the PVH of P14 male mice. (D) Schematic of our working model. Data are shown ± SEM. Statistical significance was determined using One way ANOVA (A). PVH, paraventricular nucleus of the hypothalamus; V3, third ventricle.

To confirm that *Efnb1* and *Efnb2* expressed by POMC neurons were required for the establishment of glutamatergic inputs, we used a primary co-culture cell system. PVH and ARH cells were isolated from *Pomc*-eGFP mouse cells and were maintained in culture conditions that enable the formation of neuronal networks (Figure 3A). *Efnb1* or *Efnb2* siRNA-based silencing led to an 83% and 84% decrease in *Efnb1* and *Efnb2* mRNA expression, respectively, (Figure 3B) and a 2- and 1.9-fold reduction in the number of glutamatergic terminals on POMC neurites (Figures 3C and 3D). Collectively, these results indicate that EphrinB1 and EphrinB2 positively control the amount of glutamatergic inputs into POMC neurons during the postnatal period.

**Figure 3.**
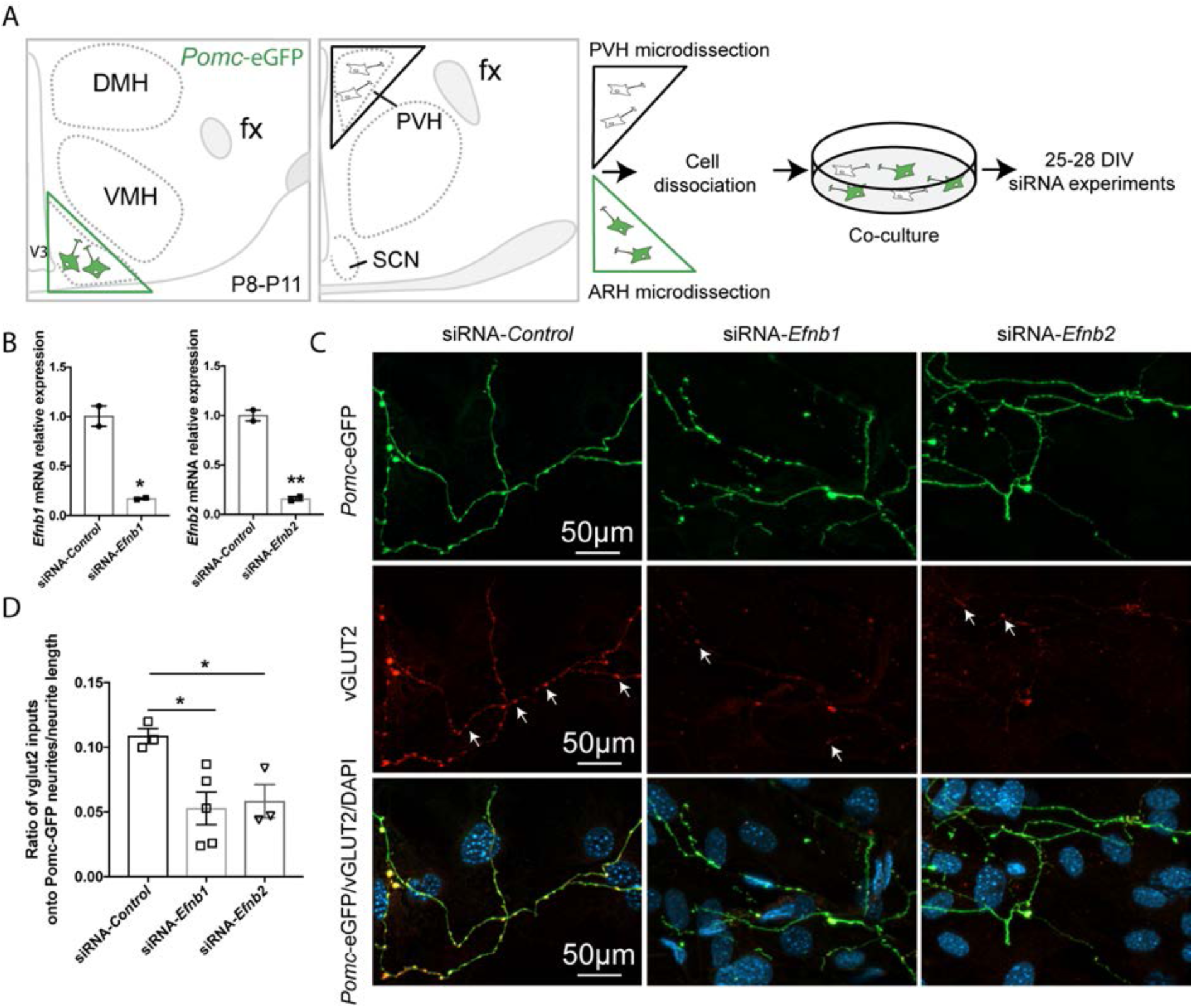
EphrinB signaling and number of glutamatergic inputs. (A) Experimental approach. ARH and PVH of *Pomc*-eGFP P8-P11 male and female mice were enzymatically dissociated and co-cultured for 25-28 days *in vitro* (DIV) before performing silencing experiments. (B) *Efnb1* and *Efnb2* mRNA relative expression upon *Efnb1* or *Efnb2* silencing or in control (siRNA-control) conditions. (C) Microphotographs showing vGLUT2-positive terminals (red) onto *Pomc*-eGFP-positive neurites (green). White arrows highlight glutamatergic inputs. (D) Quantification of glutamatergic terminals into POMC neurites. Data are shown ± SEM. Statistical significance was determined using 2-tailed Student’s t test (B, D). \**P* ≤ 0.05 versus siRNA-control (B, D); \*\**P* ≤ 0.01 *versus* siRNA-control (B). ARH, arcuate nucleus of the hypothalamus; DMH, dorsomedial nucleus of the hypothalamus; fx, fornix; PVH, paraventricular nucleus of the hypothalamus; SCN, suprachiasmatic nucleus; VMH, ventromedial nucleus of the hypothalamus; V3, third ventricle.

### Lack of *Efnb1* in POMC neurons is associated with impaired glucose homeostasis

To determine whether *Efnb1* is required for the normal development of glutamatergic synapses on POMC neurons *in vivo*, we crossed mice carrying an *Efnb1*^loxP^ allele (22) with mice expressing Cre recombinase in a *Pomc*-specific manner (23). We observed a 31% decrease in excitatory vGLUT2^+^ inputs into arcuate POMC neurons in 16-week-old *Pomc*-Cre;*Efnb1*^loxP/0^ mutant male mice (Figures 4A-C). In control mice, *Efnb1* mRNA was not detectable in the adeno-pituitary of late embryos where ACTH neurons (derived from POMC precursor) can also be found (Figure S4A); nonetheless, *Efnb1* mRNA was expressed in adult pituitary (Figure S4B and S4C). However, the deletion of *Efnb1* in POMC neurons did not affect the expression of *Efnb1* or *Efnb2* mRNA in the adult pituitary (Figures S4B and S4C).

**Figure 4.**
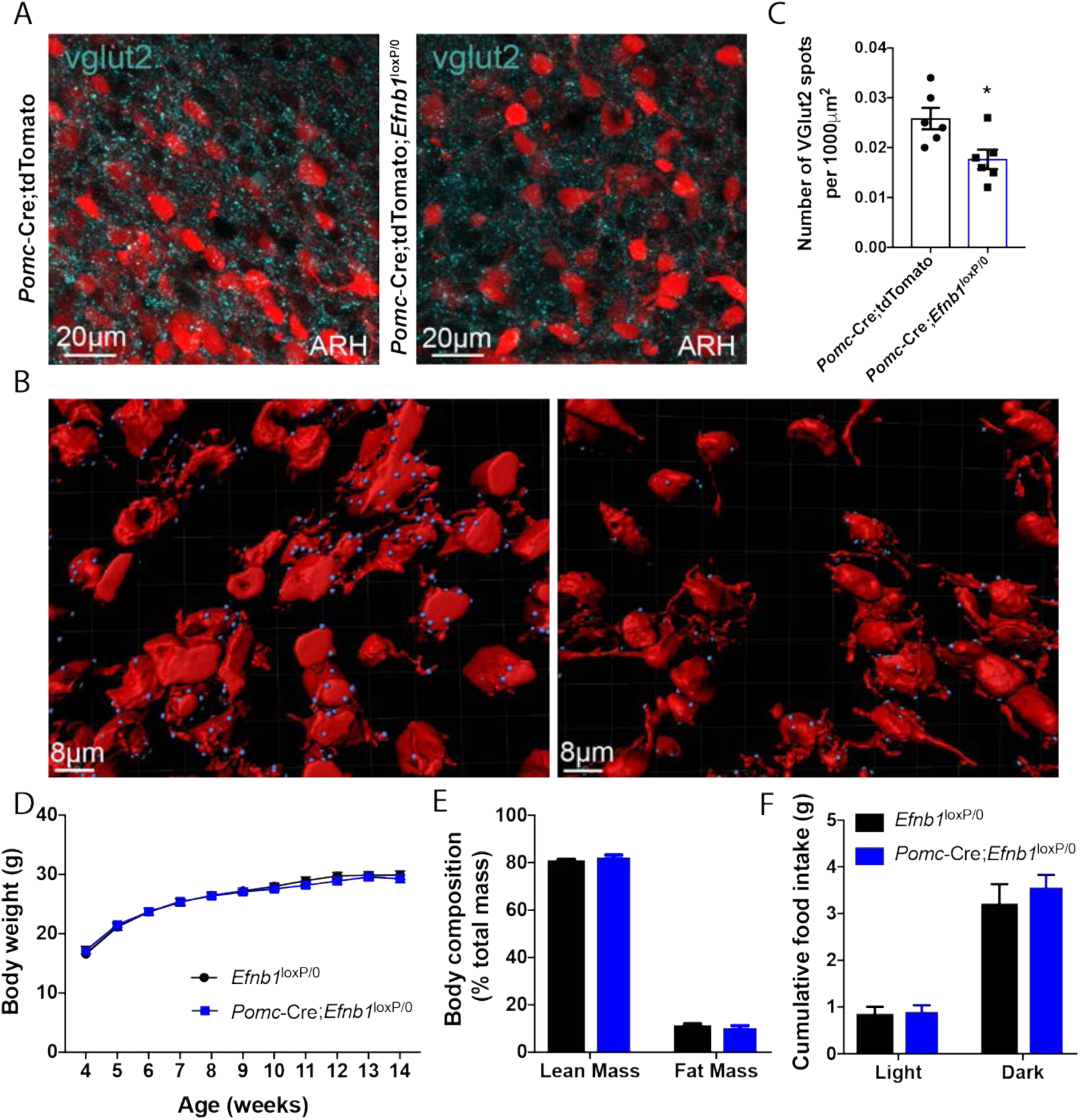
Lack of *Efnb1* in POMC neurons decreases the amount of glutamatergic inputs into POMC neurons. (A) High magnification of vGLUT2-positive terminals into POMC neurons (red) in *Pomc*-Cre;tdTomato control and *Pomc*-Cre;*Efnb1*^loxP/0^;tdTomato mutant male mice. (B) Example of 3D reconstruction (IMARIS) and quantification (C) of vGLUT2-positive inputs (blue) in direct apposition with *Pomc*-Cre;tdTomato neurons (red) (n=3/group, 2 sections/animal). (D) Post-weaning growth curve of *Efnb1*^loxP/0^ and *Pomc*-Cre;*Efnb1*^loxP/0^ male mice (n=14-15/group). (E) Body composition of 16-18 week-old male mice (n=9-10/group). (F) Cumulative food intake of 13-14 week-old *Efnb1*^loxP/0^ and *Efnb1*^loxP/0^*;Pomc*-Cre male mice (n=9-12/group). Data are shown ± SEM. Statistical significance was determined using Two Way ANOVA (D-F) and 2-tailed Student’s t test (C). \**P* ≤ 0.05 *versus Pomc*-Cre;tdTomato (C). ARH, arcuate nucleus of the hypothalamus.

Next, we examined whether the lack of *Efnb1* in POMC neurons caused disturbances in body weight and food intake. The postnatal growth curves (body weights) and body composition of *Pomc*-Cre;*Efnb1*^loxP/0^ mice were undistinguishable from *Efnb1*^loxP/0^ control male mice (Figures 4D and 4E). Consistent with these data, daily food intake was similar between the groups (Figure 4F). In addition to their fundamental role in energy balance, POMC neurons have been shown to be involved in the regulation of peripheral glucose homeostasis (5, 6). Accordingly, we also investigated the effect of genetically deleting *Efnb1* in POMC neurons on peripheral glucose homeostasis and insulin sensitivity. The basal glycemia and insulinemia were unchanged in *Pomc*-Cre;*Efnb1*^loxP/0^ mice and *Efnb1*^loxP/0^ mice (Figures 5A and 5B). *Pomc*-Cre;*Efnb1*^loxP/0^ mice displayed significantly elevated glycemia 15 to 45 minutes after a glucose challenge (Figures 5C and 5D). After 15 but not 30 minutes, glucose-stimulated insulin secretion was impaired in mutant mice, suggesting that only the cephalic phase of insulin secretion (first phase) was impacted (Figures 5E and 5F). Activation of the cholinergic parasympathetic innervation of the pancreatic islets and inhibition of the sympathetic nervous system control insulin secretion in response to hyperglycemia (24). We thus assessed cholinergic (parasympathetic) innervation of pancreatic islets in *Pomc*-Cre;*Efnb1*^loxP/0^ and *Efnb1*^loxP/0^ male mice (Figures 5G and 5H). No difference in the density of cholinergic fibers was found in islets of *Pomc*-Cre;*Efnb1*^loxP/0^ mice compared to those of *Efnb1*^loxP/0^ male mice (Figure 5H). We then measured parasympathetic nerve (vagus) activity upon glucose challenge. We observed no difference in basal firing activity between *Pomc*-Cre;*Efnb1*^loxP/0^ male mice and *Efnb1*^loxP/0^ control littermates (Figures 5I and 5J). However, whereas a glucose challenge increased firing activity by 2.5-fold in *Efnb1*^loxP/0^ control mice, no response was detected in the vagus nerve of *Pomc*-Cre;*Efnb1*^loxP/0^ mice (Figure 5J). We also performed pyruvate and insulin tolerance tests, and the results were identical in control and mutant mice (Figures S4D and S4E). Notably, similar metabolic disturbances were found in *Pomc*-Cre;*Efnb1*^loxP/loxP^ female mice (Figure S4). Together, these data show that mice lacking *Efnb1* in POMC neurons develop glucose intolerance that is associated with impaired insulin secretion and impaired parasympathetic nerve activity.

**Figure 5.**
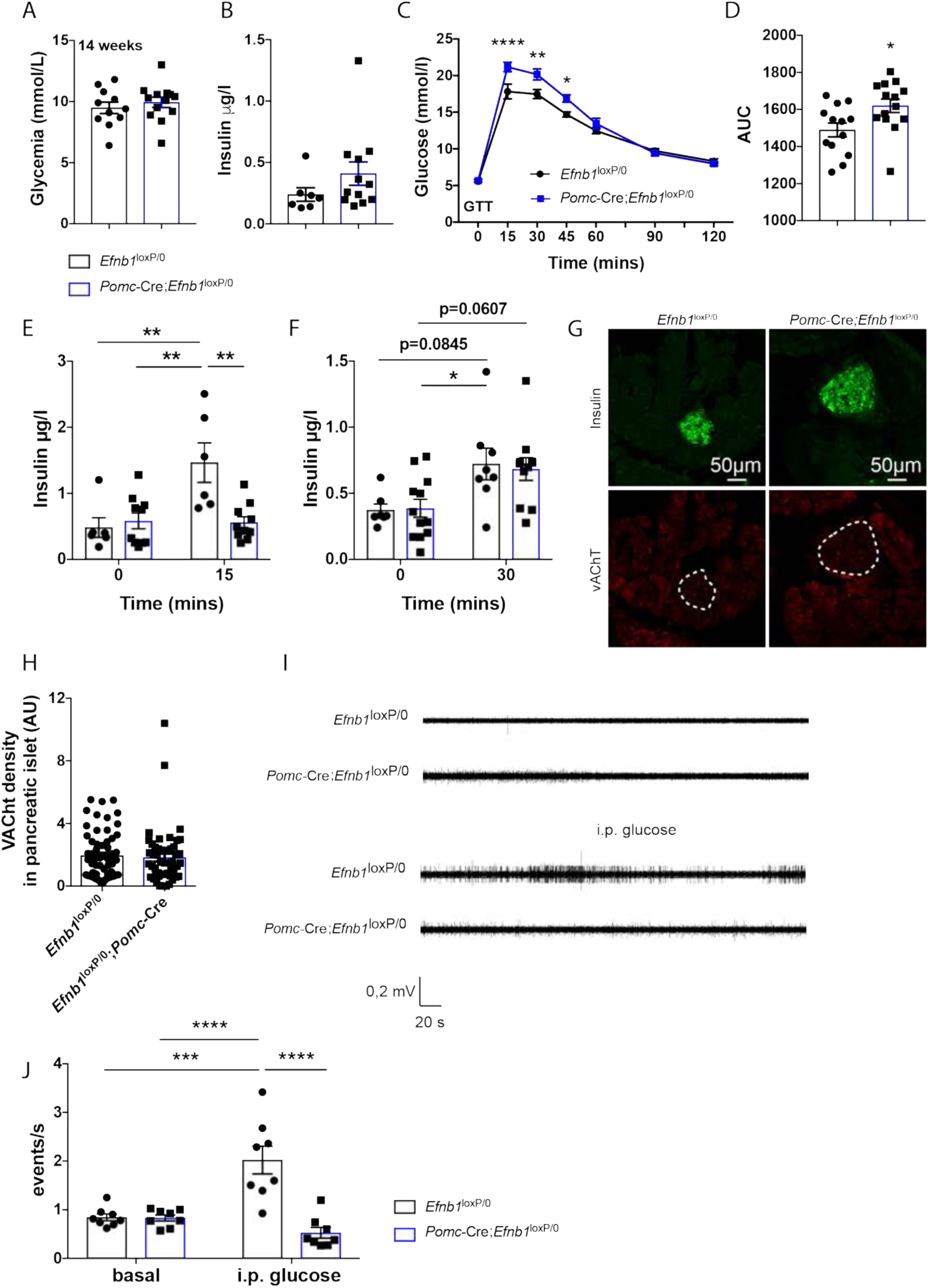
Loss of *Efnb1* in POMC neurons causes glucose intolerance in males. (A) Basal glycemia of 14 week-old male mice (n=11-13/group). (B) Basal insulinemia of 16-18 week-old male mice (n=7-12/group). (C) Glucose tolerance test of 8-9 week-old male mice (n=13-14/group). (D) Area under the curve of the glucose tolerance test. Glucose-induced insulin secretion test of 9-10 week-old male mice (n=6-12/group) 15 minutes (E) and 30 minutes (F) after glucose injection. (G) Confocal images illustrating cholinergic vAChT-positive fibers (red) in pancreatic islets labeled with insulin (green) of 16-18 week-old male mice. (H) Quantification of cholinergic fiber density in pancreatic islets (n=3 animals/group, between 16 and 27 islets per animal). Representative traces (I) and quantification (J) of parasympathetic nerve firing rate in the basal state and following i.p. glucose injection in mice (n=8/group). Data are shown ± SEM. Statistical significance was determined using Two way ANOVA (C, E, F, J) and 2-tailed Student’s t test (A, B, D, H). \**P* ≤ 0.05 *versus Efnb1*^loxP/0^ (C, D), versus *Pomc*-Cre;*Efnb1*^loxP/0^ (F); \*\**P* ≤ 0.01 *versus Efnb1*^loxP/0^ (C, E), versus *Efnb1*^loxP/0^;*Pomc*-Cre (E); \*\*\**P* ≤ 0.001 versus *Efnb1*^loxP/0^ (J), versus *Efnb1*^loxP/0^;*Pomc*-Cre (J); \*\*\*\**P* ≤ 0.0001 versus *Efnb1*^loxP/0^ (C).

### Lack of *Efnb2* in POMC neurons impairs feeding and gluconeogenesis in a sex-specific manner

To study the role of *Efnb2* in the development of glutamatergic synapses in POMC neurons, we crossed *Efnb2*^loxP^ mice (25) with *Pomc*-Cre mice. As expected, the level of *Efnb2* mRNA was significantly reduced in the ARH of *Pomc*-Cre;*Efnb2*^loxP/loxP^ mice, whereas the level of *Efnb1* mRNA was unchanged between mutant and control mice (Figures 6A and 6B). *Efnb2* mRNA was also detected in adeno-pituitary during late fetal and adult life (Figure S4A and S5A). However, no change was observed in *Efnb2* mRNA expression in the pituitaries of mice that lack *Efnb2* in their POMC cells when compared to control mice (Figure S5A).

**Figure 6.**
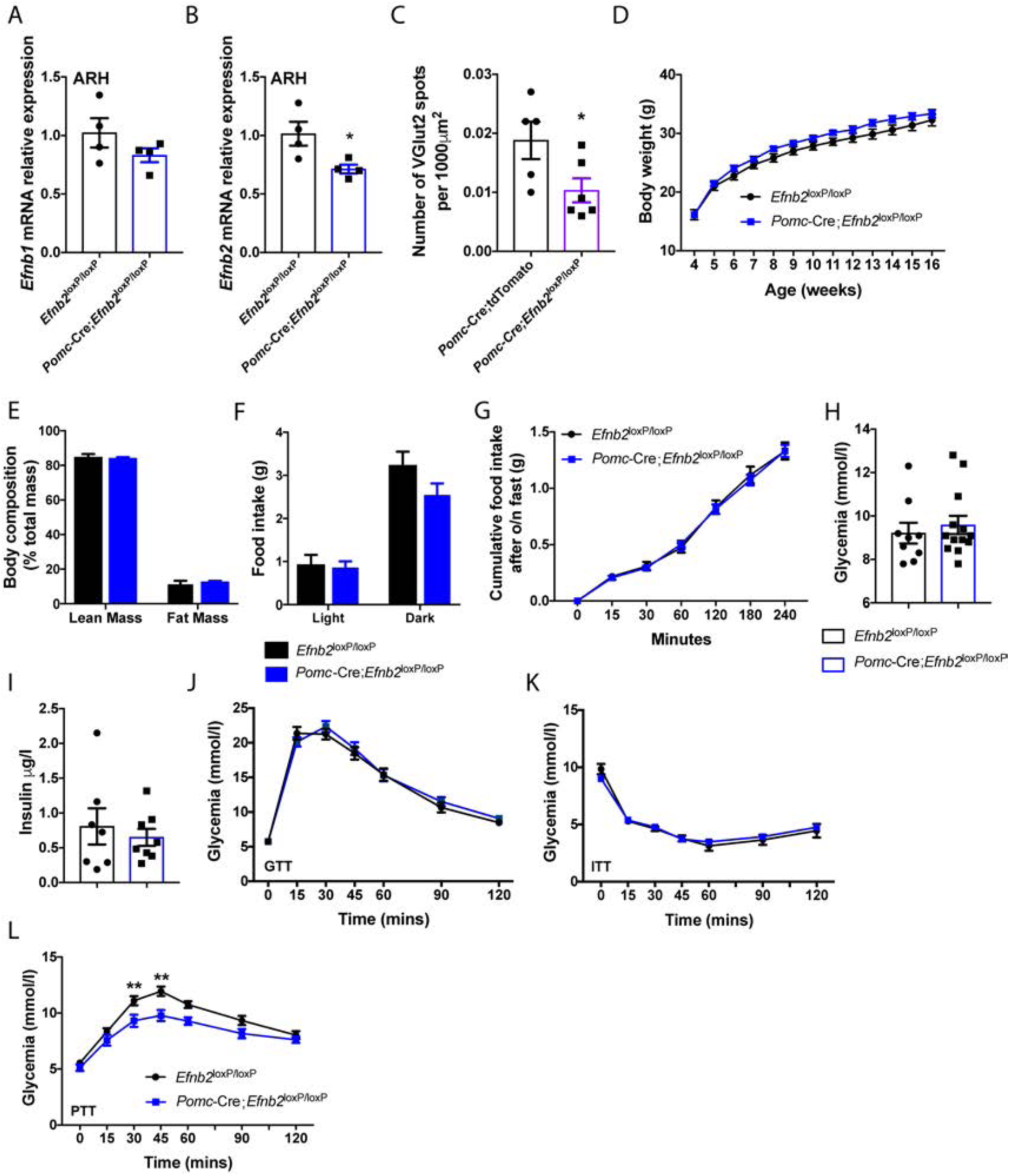
Loss of *Efnb2* in POMC neurons causes impaired gluconeogenesis in males. *Efnb1* (A) and *Efnb2* (B) mRNA relative expression in ARH of adult *Efnb2*^loxP/loxP^ and *Pomc*-Cre;*Efnb2*^loxP/loxP^ male mice (n=4/group). (C) Quantification of vGLUT2-positive inputs in direct apposition with *Pomc*-Cre;tdTomato neurons (red) in female mice (n=3/group, 2 sections/animal). (D) Post-weaning growth curve of *Efnb2*^loxP/loxP^ and *Pomc*-Cre;*Efnb2*^loxP/loxP^ male mice (n=11-14/group). (E) Body composition of 16 week-old male mice (n=7-12/group). (F) Food intake of 13-14 week-old male mice (n=6-10/group). (G) Refeeding after overnight fasting of 13-14 week-old male mice (n=13-15/group). (H) Basal glycemia of 8 week-old male mice (n=9-13/group). (I) Basal insulinemia of 16 week-old male mice (n=7-8/group). (J) Glucose tolerance test of 8-9 week-old male mice (n=11-13/group). (K) Insulin tolerance test of 14 week-old male mice (n=7-12/group). (L) Pyruvate tolerance test of 12-13 week-old male mice (n=11-13/group). Data are shown ± SEM. Statistical significance was determined using Two way ANOVA (D-G; J-L) and 2-tailed Student’s t test (A-C; H, I). \**P* ≤ 0.05 *versus Efnb2*^loxP/loxP^ (B), versus *Pomc*-Cre;tdTomato (C); \*\**P* ≤ 0.01 *versus Efnb2*^loxP/loxP^ (L).

The lack of *Efnb2* in POMC neurons affected glutamatergic inputs into POMC neurons, with a 55% decrease in the number of vGLUT2-positive terminals that were in contact with POMC neurons (Figures 6C). Physiologically, male and female mice lacking *Efnb2* in POMC neurons had normal body weight growth curves, body composition and daily food intake (Figures 6D, 6E and S5B-D). The cumulative food intake in the refeeding paradigm after overnight fasting were only comparable between control and mutant male mice (Figures 6F and 6G). Iin females, cumulative food intake after overnight fasting was significantly increased after 180 and 240 minutes in *Pomc*-Cre;*Efnb2*^loxP/loxP^ mice when compared to *Efnb2*^loxP/loxP^ mice (Figure S5E). In addition, basal glycemia, insulinemia, and glycemia levels after glucose and insulin tolerance test results were similar between the groups of male and female mice (Figures 6H-K and S5F-I).). Pyruvate tolerance tests indicated impaired gluconeogenesis only in *Pomc*-Cre;*Efnb2*^loxP/loxP^ male mice (Figure 6L and S5J).

Together, these results suggest that the lack of *Efnb2* in POMC neurons impairs gluconeogenesis in males and it impairs food intake in a refeeding paradigm in females.

## Discussion

Energy and glucose homeostasis are tightly controlled by the brain. POMC and AgRP/NPY neurons in the ARH are key regulators of these functions and respond to peripheral signals through projections to second-order neurons controlling endocrine and autonomic nervous systems. However, POMC and AgRP/NPY neurons also receive extensive inputs from a plethora of areas of the brain (18) and are thus integrated in a complex neuronal network. Although our understanding of the control of feeding behavior and glucose homeostasis has improved over the last few decades, it is still largely unknown how central circuits that regulate POMC activity and their associated functions are being assembled.

Here, we showed that there is an enrichment of EphrinB members in POMC neurons when glutamatergic inputs develop, and we described the role of ephrin signaling in the control of the amount of excitatory inputs. EphrinB1 and EphrinB2 appear to both control the number of glutamatergic terminals on POMC neurons; however, they play a distinct role in controlling energy and glucose homeostasis. These are interesting findings given that glutamatergic input pattern is impaired both in mice lacking EphrinB1 and those lacking EphrinB2 in POMC neurons, suggesting functional heterogeneity in POMC neuronal circuits.

The present study is in agreement with previous work performed in rats (20) and it shows that in mice, the amount of glutamatergic inputs into arcuate POMC neurons increases gradually after birth until weaning. During this important period of neuronal connectivity formation, we showed that POMC neurons were enriched with EphrinB1 and EphrinB2, two members of the ephrin family. These proteins enable cell to cell contacts through interaction with EphA and EphB receptors to control axon growth, synaptogenesis or synaptic plasticity. We focused our study on Ephb1 and Ephb2 receptors because their interactions with EphrinB1 and EphrinB2 are well described (21) and EphrinB members are known to play a key role in AMPA and NMDA glutamate receptor recruitment, stabilization of glutamatergic synapses, and control of the number of excitatory synapses (26, 27).

Here, we used a developmental approach that interfered with excitatory synaptic input formation to specifically assess the role of glutamatergic inputs in POMC neurons in the control of energy and glucose homeostasis. We showed that lacking EphrinB1 or EphrinB2 in POMC neurons reduces the amount of glutamatergic input into these neurons. Interestingly, these two mouse models do not have similar physiological outcomes, suggesting specificity in the establishment of glutamatergic input patterns. We first hypothesized that these differences arose from POMC heterogeneity, as distinct subsets of leptin receptor-, insulin receptor- and serotonin receptor-expressing POMC neurons are linked to functional differences; (28, 29) however, our data showed that every detected POMC neuron expressed both *Efnb1* and *Efnb2*. Thus, the differential effects of *Efnb1* and *Efnb2* inactivation on energy and glucose metabolism could stem from EphrinB favoring the formation of presynaptic inputs arising from distinct areas. Indeed, several Eph receptors can interact with EphrinB1 and EphrinB2 (21) with different affinities (30) and can also be differentially expressed in areas known to project to POMC neurons. These aspects of the study require further analysis.

The loss of *Efnb1* in POMC neurons is associated with alterations in glucose tolerance and losses of parasympathetic nerve activity and insulin secretion. These findings are consistent with previous studies, which reported that affecting either POMC signaling, circuits or POMC neuron survival leads to impaired glucose homeostasis (10–12). Surprisingly, loss of *Efnb1* in POMC neurons does not perturb food intake or body weight. Other studies have shown that ablation or inactivation of arcuate POMC neurons (12, 31) as well as genetic deficiency in POMC (32, 33) cause hyperphagia and obesity. In addition, chemogenetic stimulation of POMC neurons reduces food intake (34) as well as activation of POMC neurons projecting into the PVH (23). Notably, the aforementioned studies cannot distinguish the effects of these kinds of input from those related to the output to POMC neurons, and only a few studies have focused on the effect of synaptic inputs onto POMC neurons. Indeed, deletion of two glutamatergic NMDA receptor subunits (GluN2A and GluN2B) in POMC neurons does not lead to changes in glucose homeostasis (35). The functional effect of presynaptic input to POMC neurons can be mediated by both another NMDA subunit and through AMPA receptors (36);, both cases are in agreement with our data, in that they cannot distinguish which glutamatergic receptors are predominantly involved. In some cases, POMC functions require long or chronic chemogenetic activation (9, 12), which could reflect the recruitment of NMDA receptors alongside AMPA receptors, since Ca^2+^ entry through AMPA receptors precedes full activation of NMDA receptors (37).

POMC neurons are well known glucose-excited neurons that drive the response in maintaining normal glycemia (16, 38). Our findings may also suggest that glutamatergic innervation of POMC neurons is required for their ability to sense extracellular levels of glucose. This has been shown for NTS catecholamine neurons, which sense glucose through a presynaptic mechanism that is dependent upon, among other factors, glutamate release (39). Moreover, glucose sensing in POMC neurons requires ATP-sensitive potassium (K_ATP_) channels (16) and Kir6.2 KO mice (preventing ATP-mediated closure of KATP channels) display an absence of glutamatergic AMPA receptors (40). Finally, glucose induces glutamatergic synaptic remodeling of POMC neurons, potentially regulating the sensitivity of melanocortin system to hormonal and neuronal signals (41). Collectively these data link glucose sensing and glutamatergic release.

The loss of *Efnb2* in POMC neurons impaired gluconeogenesis in males and food intake in females in a refeeding paradigm. These findings are surprising, as chemogenetic activation and inhibition of arcuate POMC neurons repress and increase hepatic glucose production, respectively. Interestingly, a lack of ephrin expression has been shown to induce local reorganization of glutamatergic synaptic inputs (42). The lack of EphrinB2 in POMC neurons could therefore affect the number of excitatory terminals connecting to nearby AgRP neurons, which are known to suppress hepatic glucose production through insulin action (7, 8) and to promote feeding (43).

Our findings also suggest sexual dimorphism in the melanocortin system, as the lack of *Efnb2* in POMC neurons does not lead to impaired gluconeogenesis in females, but it does result in changes to refeeding after an overnight fast. The ARH has not been primarily viewed as a dimorphic structure, but recent studies showed differences between males and females in the number of ARH POMC neurons, their firing rate, the development of diet-induced obesity and the activation of STAT3 in POMC neurons (44–46).

The phenotypes we observed in mice lacking *Efnb1* or *Efnb2* can also be due to synaptic plasticity, as cells with EphrinB have been shown to control this function (27) and are found in the brains of adult animals (Allen Mouse Brain Atlas, 2004;(47). Moreover, hormones (8, 17), metabolic status, physical activity (48) and the age (49) can directly modulate the amount of glutamatergic and GABAergic synaptic inputs into POMC neurons.

In conclusion, our data show that distinct Ephrin members control the glutamatergic innervation of POMC neurons and specific functions such as glucose homeostasis or feeding. This supports the idea that POMC neuronal network is heterogeneous and that POMC neurons should not be considered as first-order neurons but have to be thought predominantly as integrators of multiple kinds of complex peripheral and central information to control energy and glucose homeostasis.

## Materials and Methods

### Experimental Model and Subject Details

#### Studies in mice

All experimental procedures were approved by the Veterinary Office of Canton de Vaud. Mice were group housed in individual cages and maintained in a temperature-controlled room with a 12hr light/dark cycle and provided *ad libitum* access to water and standard laboratory chow (Kliba Nafag). Mice were single housed only for food intake experiments.

All mice used in this study have been previously described: *Pomc*-Cre (23), ROSA-tdTomato reporter (50), *Efnb1*^loxP/loxP^ (22), *Efnb2* ^loxP/loxP^ (25), *Pomc*-eGFP (13), *Npy*-hrGFP (51). *Pomc*-Cre mice were mated to *Efnb1*^loxP/loxP^, *Efnb2* ^loxP/loxP^ to generate *Pomc*-specific *Efnb1* or *Efnb2* knockout mice. As *Efnb1* gene is carried by X chromosome, males have thus only one copy (*Efnb1*^loxP/0^).

### Method Details

#### Monosynaptic retrograde tracing

##### Virus

pAAV-syn-FLEX-splitTVA-EGFP-tTA (Addgene viral prep # 100798-AAV1; http://n2t.net/addgene:100798; RRID:Addgene_100798) and pAAV-TREtight-mTagBFP2-B19G were a gift from Ian Wickersham (Addgene viral prep # 100799-AAV1; http://n2t.net/addgene:100799; RRID:Addgene_100799).

##### Surgery

Monosynaptic retrograde tracing using rabies virus was performed as follow : Adult Pomc-Cre mice were anesthetized with a mix of xylazine ketamine. Virus were injected with a microsyringe (Hamilton, 35G) and microinjection pump (World Precisions, rate at 100nl/min). Mice receive 300nl of mixed AAV1-Syn-FLEX-splitTVA-eGFP-tTA and AAV1-TREtight-BFP2-B19G in one side of the ARH (AP : −1.4 mm; ML : −0.3 mm; DV : −5.8 mm). After 7 days, the same mice received a second injection of 300 nl of pseudotyped rabies virus EnvA-SADdG-mcherry (Salk Institute) using the same coordinates. Control mice were injected with helper virus or EnvA-SADdG-mcherry alone. One week later, mice were anesthetized and perfused with 4% PFA and frozen for brain cryosectioning.

### Cell sorting

P14 *Pomc*-Cre;tdTomato;*Npy*-hrGFP mice (n=3 for RNA-seq; n=4-8 for RT-qPCR) were microdissected under binocular loupe and enzymatically dissociated using the Papain Dissociation System (Worthington) following the manufacturer’s instructions. Fluorescence-Activated Cell Sorting (FACS) was performed using a BD FACS Aria II Cell Sorter to sort *Pomc*-tdTomato^+^, *Npy*-hrGFP^+^ and cells containg both *Pomc*-tdTomato and *Npy*-hrGFP. Non-fluorescent cells obtained from wild-type animals were used to set the gate of sorting.

### Nucleus microdissection

As already described (52), PVH and ARH nuclei were microdissected under binocular loupe from 200 µm-thick brain sections collected from P8, P10, P12, P14, P16, and P18 C57Bl/6 pups (n=2-4/age, from at least 2 litters/age) and from 16 week-old *Pomc*-Cre;*Efnb2*^loxP/loxP^ and *Efnb2*^loxP/loxP^. Microdissected nuclei were stored at −80°C until RNA extraction using Picopure RNA extraction kit (Applied Biosystems).

### RNA-sequencing

RNAs were extracted from each sorted cell population using a Picopure RNA extraction kit (Thermofischer Scientific). RNA integrity and concentration were assessed with a 2100 Bioanalyser (Agilent). 500 picograms of RNAs were reverse transcript using SMART-Seq v4 Ultra Low Input RNA (Takara) and RNA-seq librairies were prepared the Illumina Nextera XT DNA Library kit (Illumina). A sequencing depth of 98-138 millions of reads was used per library. Purity-filtered reads were adapters and quality trimmed with Cutadapt (v. 1.8, Martin 2011). Reads matching to ribosomal RNA sequences were removed with fastq_screen (v. 0.11.1). Remaining reads were further filtered for low complexity with reaper (v. 15-065, Davis et al. 2013). Reads were aligned against Mus musculus.GRCm38.92 genome using STAR (v. 2.5.3a) (53). The number of read counts per gene locus was summarized with htseq-count (v. 0.9.1) (54) using Mus musculus.GRCm38.92 gene annotation. Quality of the RNA-seq data alignment was assessed using RSeQC (v. 2.3.7) (55). Reads were also aligned to the Mus musculus.GRCm38.92 transcriptome using STAR (53) and the estimation of the isoforms abundance was computed using RSEM (v. 1.2.31) (56). Statistical analysis was performed for genes in R (R version 3.4.3). Genes with low counts were filtered out according to the rule of 1 count per million (cpm) in at least 3 samples. Library sizes were scaled using TMM normalization and log-transformed into counts per million or CPM (EdgeR package version 3.20.8) (57). Moderated t-test was used for each contrast. The adjusted p-value is computed by the Benjamini-Hochberg method, controlling for false discovery rate (FDR or adjusted p-value).

### RT-qPCR

For gene expression analyses, cDNA was generated with the high-capacity cDNA Reverse Transcription kit (Applied Biosystems). RT was performed on 70ng of RNAs for microdissected nuclei study on pups, and on 190 and 800ng, respectively, for ARH and pituitary studies in adults. RT-qPCR was performed using SyBR green mix (Applied Biosystems) and SyBR green primers (Microsynth) for *Efnb1, Efnb2, Ephb1, Ephb2, Ephb3, Ephb4, Epha4 and Epha5.* Gapdh was used for endogenous control. Primer sequences are as follow: *Efnb1 rev:* ACAGTCTCATGCTTGCCGTC*; Efnb1 fwd:* CACCCGA GCAGTTGACTACC*; Efnb2 rev:* CCTTGTCCGGGTAGAAATTTGG*; Efnb2 fwd:* GGTTTTGTGCAGAACTGCGAT*; Ephb1 rev:* CTGATGGCCTGCCAAGGTTA*; Ephb1 fwd:* CAGGCTTCACCTCCCTTCAG*; Ephb2 rev:* CTCAAACCCCCGTCTGTTACAT*; Ephb2 fwd:* CTACCCCTCATCGTTGGCTC*; Ephb3 rev:* GAGCTGAGTGTCA GACCTGC*; Ephb3 fwd:* GACTGCAGAAGATCTGCTAAGGA*; Ephb4 rev:* TCTGCGCCCTTCTCATGATACT*; Ephb4 fwd:* TTTCTTTC TCCTGCAGTGCCT*; Epha4 rev:* AGTTCGCAGCCAGTTGTTCT*; Epha4 fwd:* CTGGAAGG AGGGTGGGAGG*; Epha5 rev:* ATTCCATTGGGGCGATCTGG*; Epha5 fwd:* GGTACCTGCCAAGCTCCTTC; *Gapdh rev:* AAGATGGTGATGGGCTTCCC*; Gapdh fwd:* CTCCACTCACGGCAAATTCA.

All assays were performed using an Applied Biosystems 7500 Fast real-time PCR system. Calculations were performed by comparative method (2-ΔΔCT).

### *In vitro* assay

P8 to P11 *Pomc*-eGFP male and female were used for the *in vitro* study. After brain collection, PVH- and ARH-containing areas were microdissected under binocular loupe. Cells were dissociated using the Papain Dissociation System (Worthington) following the manufacturer’s instructions. After dissociation, 40’000 cells from a mix of PVH and ARH cells (ratio 1:1.2 to 1.4) were plated in poly-L-lysine coated coverslip in 24-well plates containing Neurobasal medium supplemented with 1X B27, 200mM glutamine, 1X penicillin/streptomycin. One hour later, medium was changed and plates were incubated for a week at 37°C under 5% CO2 before weekly changing the medium. After 25-28 days *in vitro* (DIV), cultures were exposed to 50nM of ON-Target plus Mouse *Efnb1* siRNA (J-051250-10-0002, #13642, Dharmacon), ON-Target plus Mouse *Efnb2* siRNA (J-051250-09-0002, #13641, Dharmacon) or ON-Target plus Non-targeting Pool control siRNA (D-001810-10-05, Dharmacon) and lipofectamine 2000 (Thermo scientific). After 3 days, cells were either fixed for 30min with 4% paraformaldehyde in PBS or collected for RNA extraction and RT-qPCR to assess siRNA-silencing efficiency. Cells were then incubated in rabbit anti-vGLUT2 (1:500, Synaptic Systems) and chicken anti-GFP (1:500, Abcam) for 2 days at 4°C. The primary antibodies were visualized with Alexa goat anti-rabbit 568 and Alexa goat anti-chicken 488. Coverslips were then mounted with DAPI fluoromount. Number of glutamatergic inputs were manually quantified using Image J.

### *In situ* hybridization (RNAscope)

P14 male C57Bl/6, *Pomc*-eGFP, and 14-week old *Pomc*-Cre male mice (for monosynaptic retrograde tracing) were perfused with 4% PFA. WT embryos at embryonic day E17 were collected and immerged in 4% PFA overnight. On 20 µm-thick brain or embryo coronal cryosections, *in situ* hybridization for *Efnb1* (cat # 526761, cat # 526761-C2), *Efnb2* (cat # 477671), *Ephb1* (cat # 567571-C3), *Ephb2* (cat # 447611-C3), *Slc17a6 (vglut2)* (cat # 319171) was processed using RNAscope probes and RNAscope Fluorescent Multiplex Detection Reagents (Advanced Cell Diagnostics) following manufacturer’s instructions.

### Immunohistochemistry

P6, P14, and P22 *Pomc*-Cre;tdTomato male mice (n=2-3 animals/age), 16-18 week-old *Pomc*-Cre;tdTomato;*Efnb1*^loxP/0^, *Pomc*-Cre;tdTomato, *Pomc*-Cre;tdTomato;*Efnb2*^loxP/loxP^, and *Pomc*-Cre;tdTomato;*Efnb1*^loxP/loxP^, *Pomc*-Cre;tdTomato;*Efnb2*^loxP/loxP^*, Pomc*-Cre;tdTomato female mice were transcardially perfused with 4% PFA (n=3/group). 20 µm-thick brain and pancreas sections were processed for immunofluorescence using standard procedures (10,52,58). The primary antibodies used for IHC were as follows: rabbit anti-vGLUT2 (1:500, Synaptic Systems), rabbit anti-VAChT (1:500, Synaptic Systems), guinea-pig anti-insulin (1:500, Abcam). Primary antibody was visualized with Alexa anti-rabbit 647 and 568 and Alexa anti-guinea-pig 488.

### Images analyses

To quantitatively analyze cholinergic (VAChT-positive) fibers in pancreatic cells, between 16 and 27 pancreatic islets per animal were imaged using a Zeiss LSM 710 confocal system equipped with a 20X objective. Each image was binarized to isolate labeled fibers from the background and to compensate for differences in fluorescence intensity. The integrated intensity, which reflects the total number of pixels in the binarized image, was then calculated for each islet. This procedure was conducted for each image. Image analysis was performed using Image J analysis software (NIH).

To quantitatively analyze glutamatergic innervation of POMC neurons, adjacent image planes were collected in lateral part of the ARH through the z-axis using a Zeiss LSM 710 confocal system at a frequency of 0.25 µm through the entire thickness of the ARH. Three-dimensional reconstructions of the image volumes were then prepared using Imaris 9.3.1 visualization software. The number of glutamatergic inputs into POMC neurons was quantified. Each putative glutamatergic input was defined as a spot, and we quantified the number of glutamatergic spots that contacted *Pomc*-Cre;tdtomato+.

### Physiological measures

*Pomc*-Cre;tdTomato;*Efnb1*^loxP/0^, *Efnb1*^loxP/0^ (n = 14-16 per group), *Pomc*-Cre;tdTomato;*Efnb2*^loxP/loxP^, *Efnb2*^loxP/loxP^ (n = 11-14 per group) male and *Pomc*-Cre;tdTomato;*Efnb1*^loxP/loxP^, *Efnb1*^loxP/loxP^ (n = 8-10 per group), *Pomc*-Cre;tdTomato;*Efnb2*^loxP/loxP^, *Efnb2*^loxP/loxP^ female mice (n = 19 per group) were weighed every week from 3 weeks (weaning) to 16 weeks using an analytical balance. To measure food consumption, 13-14-week-old *Pomc*-Cre;tdTomato;*Efnb1*^loxP/0^, *Efnb1*^loxP/0^ (n = 9-12 per group), *Pomc*-Cre;tdTomato;*Efnb2*^loxP/loxP^, *Efnb2*^loxP/loxP^ (n = 6-10 per group) male mice and *Pomc*-Cre;tdTomato;*Efnb2*^loxP/loxP^, *Efnb2*^loxP/loxP^ (n = 8 per group) female mice were housed individually in BioDAQ cages (Research Diets), and, after at least 2 days of acclimation, food intake was assessed on 2 consecutive days. The means obtained on these 2 days were used for analyses. Body composition analysis (fat/lean mass) was performed in 16-week-old *Pomc*-Cre;tdTomato;*Efnb1*^loxP/0^, *Efnb1*^loxP/0^ (n = 9-10 per group), *Pomc*-Cre;tdTomato;*Efnb2*^loxP/loxP^, *Efnb2*^loxP/loxP^ (n = 7-12 per group) male mice and *Pomc*-Cre;tdTomato;*Efnb2*^loxP/loxP^, *Efnb2*^loxP/loxP^ (n = 6 per group) female mice using NMR (Echo MRI). Glucose (GTT), insulin (ITT) and pyruvate (PTT) tolerance tests were conducted in 8- to 12 week-old *Pomc*-Cre;tdTomato;*Efnb1*^loxP/0^, *Efnb1*^loxP/0^ (n = 9-14 per group), *Pomc*-Cre;tdTomato;*Efnb2*^loxP/loxP^, *Efnb2*^loxP/loxP^ (n = 8-13 per group) male mice and *Pomc*-Cre;tdTomato;*Efnb2*^loxP/loxP^, *Efnb2*^loxP/loxP^ (n = 8-14 per group) female mice through i.p. injection of glucose (2 mg/g body weight), insulin (0.5 U/kg body weight) or sodium pyruvate (2 mg/g body weight) after overnight fasting (15h-16h, GTT and PTT) or 5-6 h of fasting (ITT). Blood glucose levels were measured at 0, 15, 30, 45, 60, 90, and 120 min post-injection. Glycemia was measured using a glucometer (Bayer).

Glucose-stimulated insulin secretion tests were also performed at 9-10 weeks of age, through the i.p. administration of glucose (2 mg/kg body weight, n = 6-12 per group) after 15-16h overnight fasting. Blood samples were collected 0, 15 min and 0 and 30 min after glucose injection on 2 distinct cohortes. Serum insulin levels were then measured using an insulin ELISA kit (Mercodia). Basal insulinemia was measured on 16-18 week-old mice (n=6-12/group) using an insulin ELISA kit (Mercodia).Basal glycemia was measured the morning on fed mice using a glucometer (Bayer).

### Vagus nerve activity recording

The firing rate of the thoracic branch of the vagal nerve along the carotid artery was recorded as previously described (59–61) on 14-15 week-old *Pomc*-Cre;tdTomato;*Efnb1*^loxP/0^, *Efnb1*^loxP/0^ (n = 8/ group). Unipolar nerve activity was recorded continuously under isoflurane anesthesia (30 min during basal condition and 30 min after i.p. glucose at a dose of 2g/kg) using the LabChart 8 software (AD Instrument, Oxford, UK). Data were digitized with PowerLab 16/35 (AD Instrument, Oxford, UK). Signals were amplified 10^5^ times and filtered using 200/1000 Hz band pass filter. Firing rate analysis was performed using LabChart 8. Data were analyzed on 6 min at the end of the basal recording and for the same duration 15 min after glucose i.p. injection.

### Quantification and statistical analysis

All values were represented as the mean ± SEM. Numbers for every experiment are found in the relevant part of the STAR Methods. Statistical analyses were conducted using GraphPad Prism (version 7). Statistical significance was determined using unpaired 2-tailed Student’s t test, 1-way ANOVA followed by Tukey’s post hoc test, 2-way ANOVA followed by Sidak’s post hoc test when appropriate. P ≤ 0.05 was considered statistically significant.

## Authors Contributions

S.C. conceived the project, designed the experiments, analyzed the data and wrote the manuscript with inputs from A.P and B.T authors. S.C. performed all experiments except the RNA-sequencing (GTF facility), RT-qPCR, body weight, body composition, glycemia, (M.G.), vagus nerve recording (A.P.).

## Acknowledgments

We thank the CIG Genomic Technologies Facility (GTF) for RNA-sequencing experiments and analyses, the EPFL Flow Cytometry Core, the UNIL Flow Cytometry Facilities for cell sorting, and the CIG Animal Facility for their assistance with animal husbandry. We are also grateful to Marc Lanzillo with his assistance with animal genotyping. This work was supported by the Swiss National Science Foundation grant (PZ00P3_167934/1) and Novartis grant (19B145).

---

**Figure S1.**
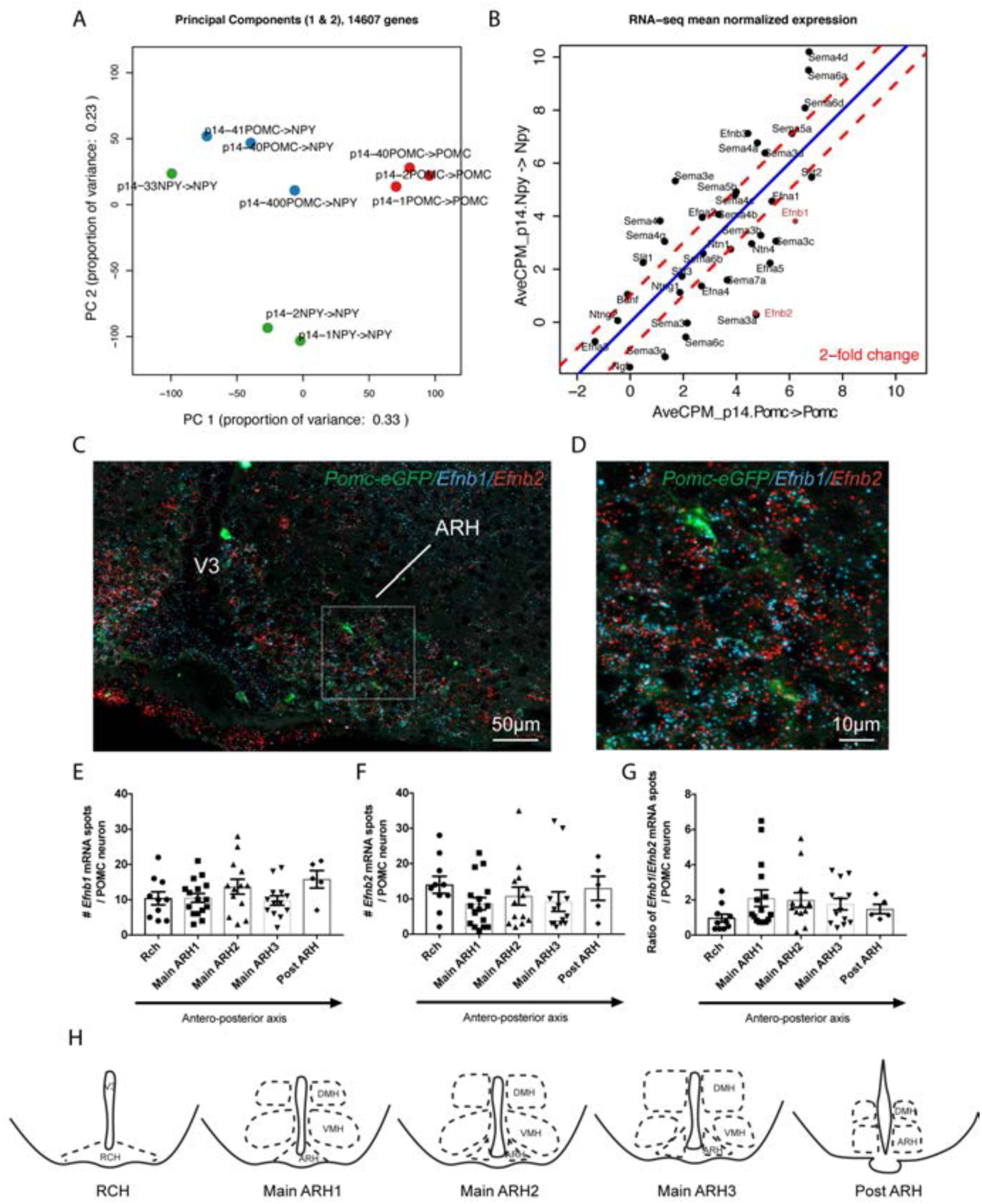
*Efnb1* and *Efnb2* mRNA are enriched in POMC neurons. (A) Principal component analysis made on 14607 genes. (B) Scatterplots comparing the expression of individual genes between Pomc->Pomc and Npy->Npy neuronal population at P14. (C) Microphotographs showing *Efnb1* (blue) and *Efnb2* (red) mRNA spots in POMC-GFP^+^ (green) neurons in the ARH of P14 male mice. (D) High magnification of the inset shown in (C). Quantification of the number of *Efnb1* mRNA spots (E), *Efnb2* mRNA spots (F) or *Efnb1*/*Efnb2* mRNA spot ratio (G) in POMC-eGFP neurons in the entire thickness of the ARH of P14 male animals (n=2 animals). (H) Schematic illustrating the subdivisions of the ARH used for the quantification in E, F and G. Data are shown ± SEM. Statistical significance was determined using One way ANOVA (E-G). ARH, arcuate nucleus of the hypothalamus; DMH, dorsomedial nucleus of the hypothalamus; VMH, ventromedial nucleus of the hypothalamus; V3, third ventricle; RCH, retrochiasmatic area.

**Figure S2.**
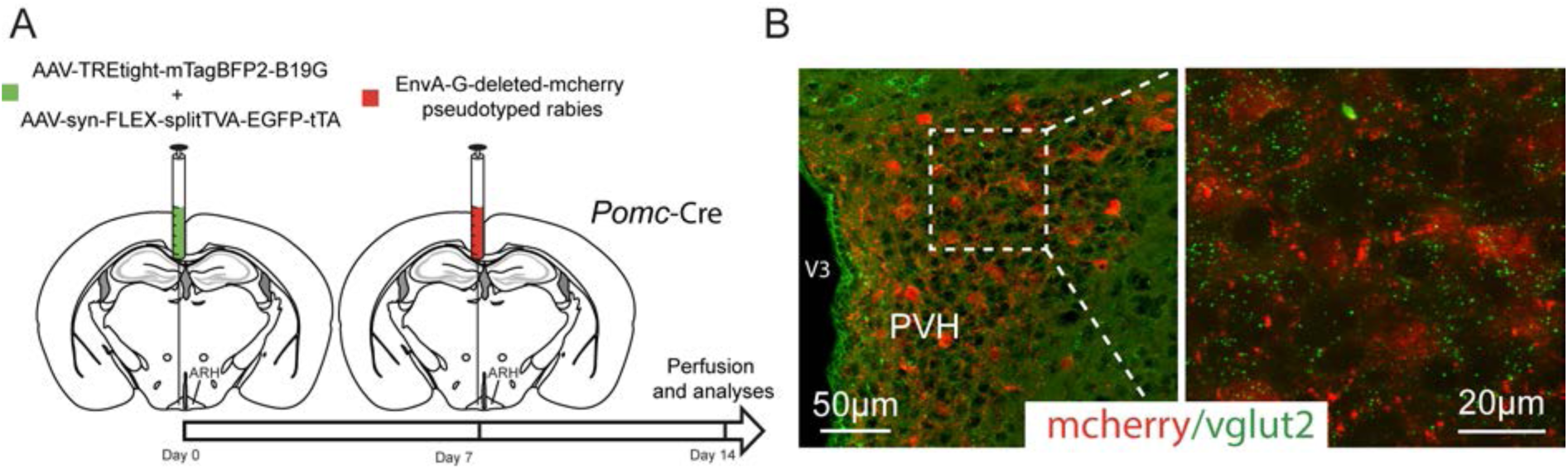
Glutamatergic PVH neurons project into POMC neurons of the ARH. (A) Experimental approach. A mix of AAV-TREtight-mTagBFP2-B19G and AAV-syn-FLEX-splitTVA-EGFP-tTA was injected at day 0 in the ARH of 12 week-old *Pomc*-Cre male mice. Seven days later, mice received injection of EnvA-G-deleted-mcherry pseudotyped rabies. Animals were perfused one week later for further analyses. (B) Photomicrographs showing the co-localization of mcherry-positive cells (POMC inputs) with glutamatergic neurons of the PVH (*vglut2* mRNA spots in green). ARH, arcuate nucleus of the hypothalamus; PVH, paraventricular nucleus of the hypothalamus; V3, third ventricle.

**Figure S3.**
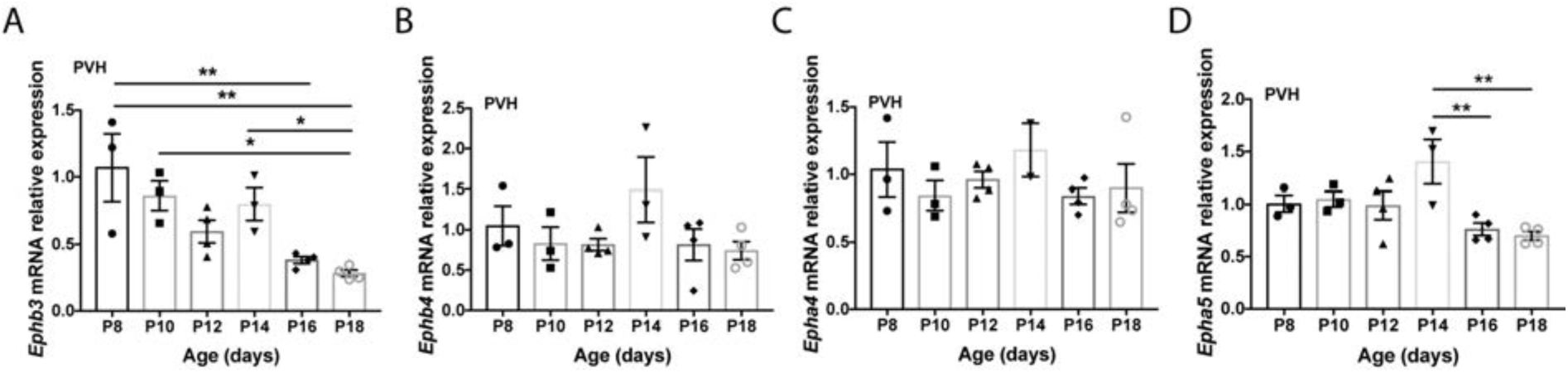
Eph receptors are expressed in PVH during postnatal development. Quantification of *Ephb3* (A), *Ephb4* (B), *Epha4* (C) and *Epha5* (D) mRNA relative expression in PVH of P8, P10, P12, P14, P16 and P18 male pups (n=2-4 pups/age). Data are shown ± SEM. Statistical significance was determined using One way ANOVA (A-D). \**P* ≤ 0.05 *versus* P12 (A), versus P14 (A), \*\**P* ≤ 0.01 *versus* P8 (A), versus P14 (D).

**Figure S4.**
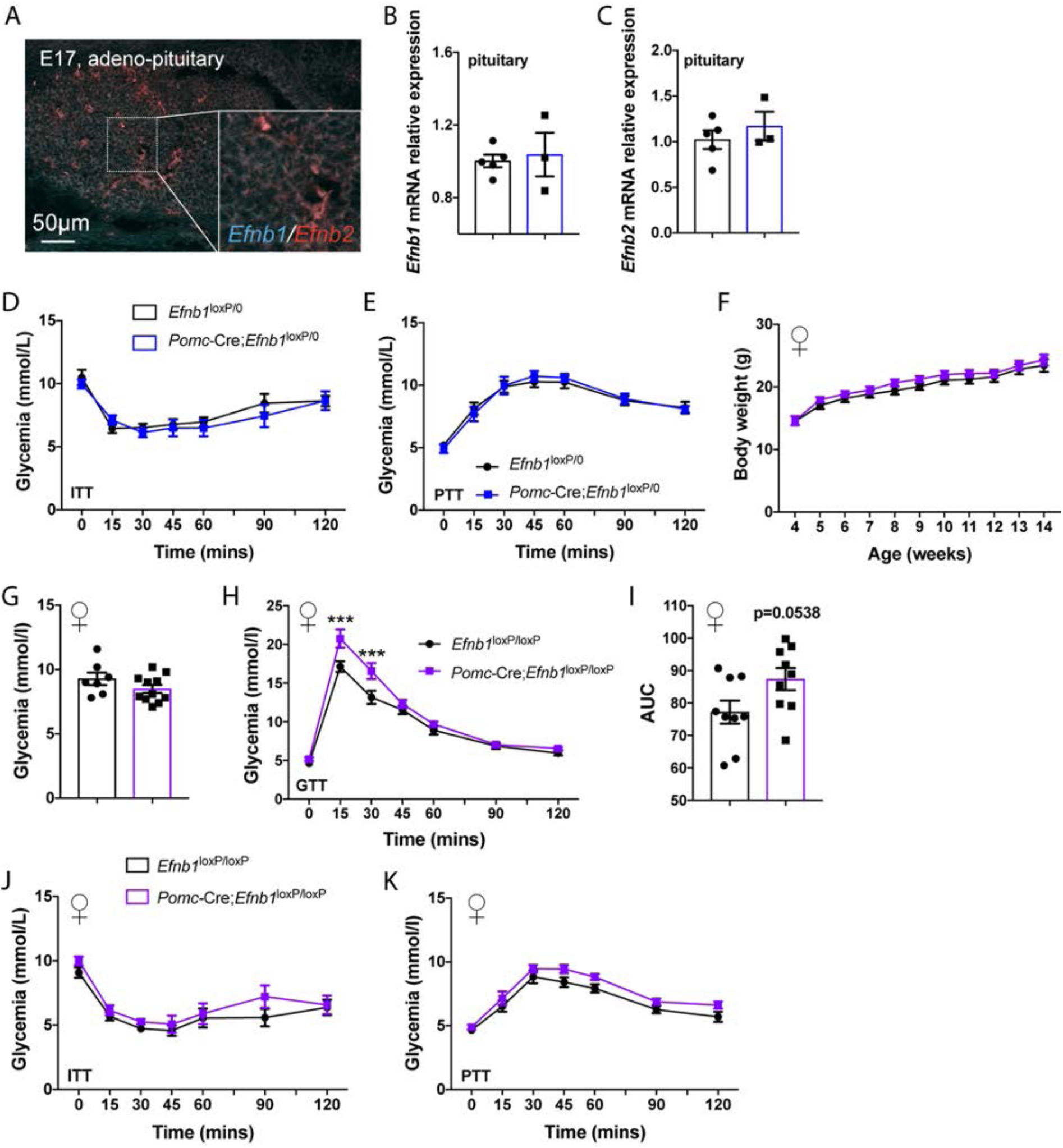
Loss of *Efnb1* in POMC neurons causes glucose intolerance in females. (A) Microscope image illustrating the expression of *Efnb1* (blue) and *Efnb2* (red) mRNA in the adeno-pituitary of E17 embryo. (B) *Efnb1* and *Efnb2* mRNA relative expression in the pituitary of *Efnb1*^loxP/0^ and *Pomc*-Cre;*Efnb1*^loxP/0^ 16-week old male mice (n=3-5/group). (D) Insulin tolerance test of 14 week-old male mice (n=10-13/group). (E) Pyruvate tolerance test of 13 week-old male mice (n=9-10/group). (F) Post-weaning growth curve of *Efnb1*^loxP/loxP^ and *Pomc*-Cre;*Efnb1*^loxP/loxP^ female mice (n=8-11/group). (G) Basal glycemia of 8 week-old female mice (n=7-11/group). (H) Glucose tolerance test of 8-9 week-old female mice (n=9/group). (I) Area under the curve of GTT experiment. (J) Insulin tolerance test of 14 week-old female mice (n=9/group). (K) Pyruvate tolerance test of 13 week-old female mice (n=7-8/group). Data are shown ± SEM. Statistical significance was determined using Two way ANOVA (D-F, H, J, K) and 2-tailed Student’s t test (B, C, G, I). \*\*\**P* ≤ 0.001 *versus Efnb1*^loxP/loxP^ (H).

**Figure S5.**
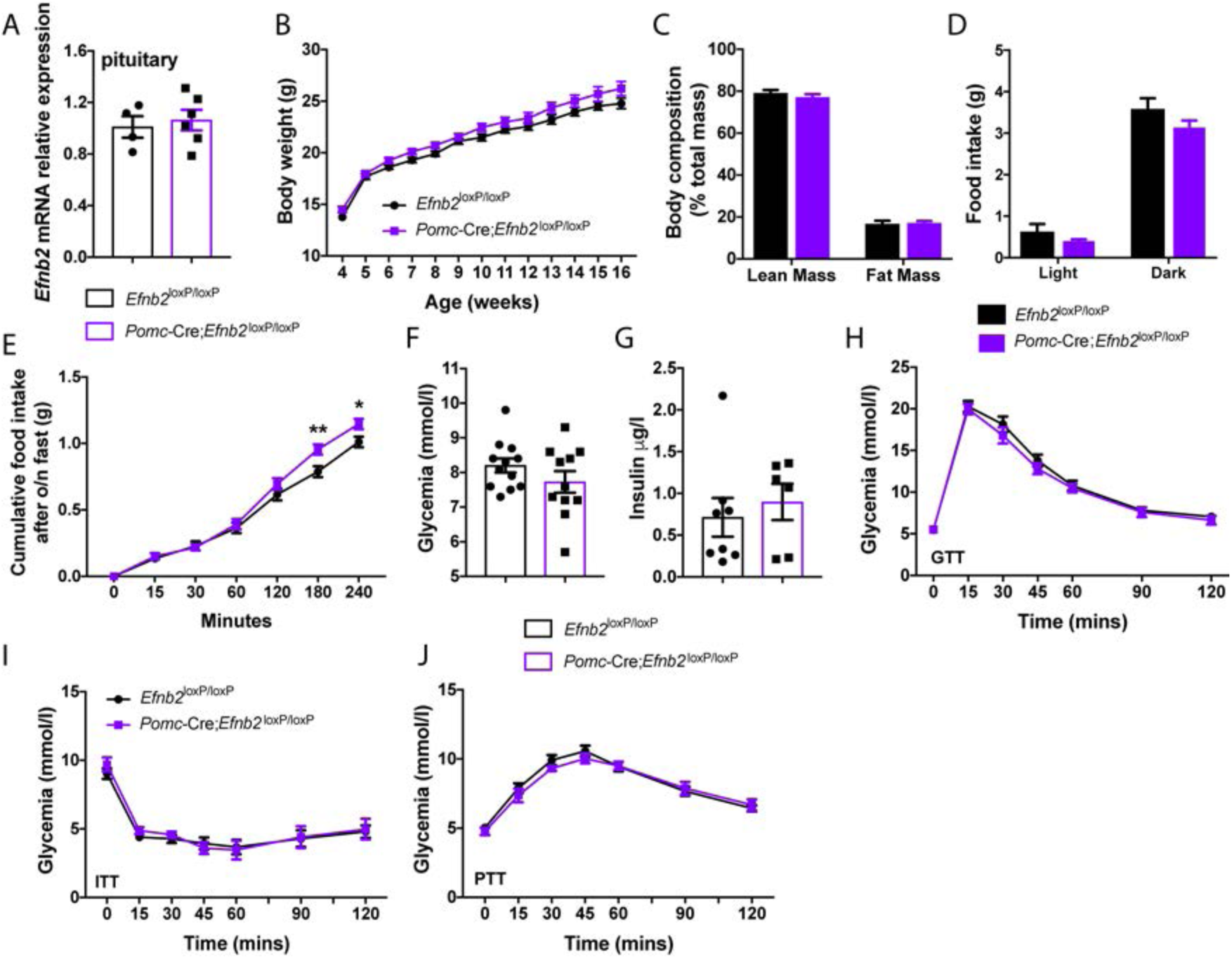
Loss of *Efnb2* in POMC neurons causes impaired refeeding after overnight fast in females. (A) *Efnb2* mRNA relative expression in the pituitary of *Efnb2*^loxP/loxP^ and *Pomc*-Cre;*Efnb2*^loxP/loxP^ male mice (n=4-6/group). (B) Post-weaning growth curve of *Efnb2*^loxP/loxP^ and *Pomc*-Cre;*Efnb2*^loxP/loxP^ female mice (n=19/group). (C) Body composition of 16 week-old female mice (n=6/group). (D) Food intake of 13-14 week-old female mice (n=8/group). (E) Refeeding after overnight fasting of 13-14 week-old female mice (n=11-15/group). (F) Basal glycemia of 8 week-old female mice (n=11-12/group). (G) Basal insulinemia of 16 week-old female mice (n=6-8/group). (H) Glucose tolerance test of 8-10 week-old female mice (n=13-14/group). (I) Insulin tolerance test of 14 week-old female mice (n=8-10/group). (J) Pyruvate tolerance test of 12-13 week-old female mice (n=8-14/group). Data are shown ± SEM. Statistical significance was determined using Two way ANOVA (B-E, H-J) and 2-tailed Student’s t test (A, F, G). \**P* ≤ 0.05 *versus Efnb2*^loxP/loxP^ (E); \*\**P* ≤ 0.01 *versus Efnb2*^loxP/loxP^ (E).

**Table S1. List of putative genes involved in synapse formation and axon guidance.**

## References

1. Gautron L, Elmquist JK, Williams KW. Neural control of energy balance: translating circuits to therapies. Cell. 2015 Mar 26;161(1):133–45.

2. van der Klaauw AA, Farooqi IS. The hunger genes: pathways to obesity. Cell. 2015 Mar 26;161(1):119–32.

3. Timper K, Brüning JC. Hypothalamic circuits regulating appetite and energy homeostasis: pathways to obesity. Disease Models & Mechanisms. 2017 Jun 1;10(6):679–89.

4. Zhan C. POMC Neurons: Feeding, Energy Metabolism, and Beyond. In: Wu Q, Zheng R, editors. Neural Regulation of Metabolism [Internet]. Singapore: Springer Singapore; 2018 [cited 2019 Mar 20]. p. 17–29. (Advances in Experimental Medicine and Biology). Available from: https://doi.org/10.1007/978-981-13-1286-1_2

5. Pozo M, Claret M. Hypothalamic Control of Systemic Glucose Homeostasis: The Pancreas Connection. Trends in Endocrinology & Metabolism. 2018 Aug 1;29(8):581– 94.

6. Ruud J, Steculorum SM, Brüning JC. Neuronal control of peripheral insulin sensitivity and glucose metabolism. Nat Commun. 2017 May 4;8:15259.

7. Könner AC, Janoschek R, Plum L, Jordan SD, Rother E, Ma X, et al. Insulin action in AgRP-expressing neurons is required for suppression of hepatic glucose production. Cell Metab. 2007 Jun;5(6):438–49.

8. Lin HV, Plum L, Ono H, Gutiérrez-Juárez R, Shanabrough M, Borok E, et al. Divergent regulation of energy expenditure and hepatic glucose production by insulin receptor in agouti-related protein and POMC neurons. Diabetes. 2010 Feb;59(2):337–46.

9. Dodd GT, Michael NJ, Lee-Young RS, Mangiafico SP, Pryor JT, Munder AC, et al. Insulin regulates POMC neuronal plasticity to control glucose metabolism. eLife [Internet]. 2018 [cited 2019 Nov 26];7. Available from: https://www.ncbi.nlm.nih.gov/pmc/articles/PMC6170188/

10. Croizier S, Park S, Maillard J, Bouret SG. Central Dicer-miR-103/107 controls developmental switch of POMC progenitors into NPY neurons and impacts glucose homeostasis. Elife. 2018 Oct 12;7.

11. Vogt MC, Paeger L, Hess S, Steculorum SM, Awazawa M, Hampel B, et al. Neonatal insulin action impairs hypothalamic neurocircuit formation in response to maternal high-fat feeding. Cell. 2014 Jan 30;156(3):495–509.

12. Zhan C, Zhou J, Feng Q, Zhang J-E, Lin S, Bao J, et al. Acute and long-term suppression of feeding behavior by POMC neurons in the brainstem and hypothalamus, respectively. J Neurosci. 2013 Feb 20;33(8):3624–32.

13. Cowley MA, Smart JL, Rubinstein M, Cerdán MG, Diano S, Horvath TL, et al. Leptin activates anorexigenic POMC neurons through a neural network in the arcuate nucleus. Nature. 2001 May 24;411(6836):480–4.

14. Cowley MA, Smith RG, Diano S, Tschöp M, Pronchuk N, Grove KL, et al. The distribution and mechanism of action of ghrelin in the CNS demonstrates a novel hypothalamic circuit regulating energy homeostasis. Neuron. 2003 Feb 20;37(4):649– 61.

15. Hill JW, Elias CF, Fukuda M, Williams KW, Berglund ED, Holland WL, et al. Direct Insulin and Leptin Action in Pro-opiomelanocortin Neurons is Required for Normal Glucose Homeostasis and Fertility. Cell Metab. 2010 Apr 7;11(4):286–97.

16. Parton LE, Ye CP, Coppari R, Enriori PJ, Choi B, Zhang C-Y, et al. Glucose sensing by POMC neurons regulates glucose homeostasis and is impaired in obesity. Nature. 2007 Sep 13;449(7159):228–32.

17. Vong L, Ye C, Yang Z, Choi B, Chua S, Lowell BB. Leptin Action on GABAergic Neurons Prevents Obesity and Reduces Inhibitory Tone to POMC Neurons. Neuron. 2011 Jul 14;71(1):142–54.

18. Wang D, He X, Zhao Z, Feng Q, Lin R, Sun Y, et al. Whole-brain mapping of the direct inputs and axonal projections of POMC and AgRP neurons. Front Neuroanat [Internet]. 2015 Mar 27 [cited 2018 Jul 30];9. Available from: https://www.ncbi.nlm.nih.gov/pmc/articles/PMC4375998/

19. Pinto S, Roseberry AG, Liu H, Diano S, Shanabrough M, Cai X, et al. Rapid rewiring of arcuate nucleus feeding circuits by leptin. Science. 2004 Apr 2;304(5667):110–5.

20. Matsumoto A, Arai Y. Developmental changes in synaptic formation in the hypothalamic arcuate nucleus of female rats. Cell Tissue Res. 1976 Jun 14;169(2):143– 56.

21. Gaitanos, Dudanova I, Sakkou M, Klein R, Paixão S. Receptor Tyrosine Kinases: Family and Subfamilies. Springer, Cham; 2015.

22. Davy A, Aubin J, Soriano P. Ephrin-B1 forward and reverse signaling are required during mouse development. Genes Dev. 2004 Mar 1;18(5):572–83.

23. Balthasar N, Dalgaard LT, Lee CE, Yu J, Funahashi H, Williams T, et al. Divergence of melanocortin pathways in the control of food intake and energy expenditure. Cell. 2005 Nov 4;123(3):493–505.

24. Steinbusch L, Labouèbe G, Thorens B. Brain glucose sensing in homeostatic and hedonic regulation. Trends Endocrinol Metab. 2015 Sep;26(9):455–66.

25. Grunwald IC, Korte M, Adelmann G, Plueck A, Kullander K, Adams RH, et al. Hippocampal plasticity requires postsynaptic ephrinBs. Nat Neurosci. 2004 Jan;7(1):33–40.

26. Henderson N, Dalva MB. EphBs and ephrin-Bs: Trans-synaptic organizers of synapse development and function. Molecular and Cellular Neuroscience [Internet]. 2018 Jul 19 [cited 2018 Jul 30]; Available from: http://www.sciencedirect.com/science/article/pii/S1044743118301453

27. Hruska M, Dalva MB. Ephrin regulation of synapse formation, function and plasticity. Mol Cell Neurosci. 2012 May;50(1):35–44.

28. Sohn J-W, Xu Y, Jones JE, Wickman K, Williams KW, Elmquist JK. Serotonin 2C Receptor Activates a Distinct Population of Arcuate Pro-opiomelanocortin Neurons via TRPC Channels. Neuron. 2011 Aug 11;71(3):488–97.

29. Williams KW, Margatho LO, Lee CE, Choi M, Lee S, Scott MM, et al. Segregation of acute leptin and insulin effects in distinct populations of arcuate proopiomelanocortin neurons. J Neurosci. 2010 Feb 17;30(7):2472–9.

30. Blits-Huizinga CT, Nelersa CM, Malhotra A, Liebl DJ. Ephrins and their receptors: binding versus biology. IUBMB Life. 2004 May;56(5):257–65.

31. Xu AW, Kaelin CB, Morton GJ, Ogimoto K, Stanhope K, Graham J, et al. Effects of hypothalamic neurodegeneration on energy balance. PLoS Biol. 2005 Dec;3(12):e415.

32. Krude H, Biebermann H, Luck W, Horn R, Brabant G, Grüters A. Severe early-onset obesity, adrenal insufficiency and red hair pigmentation caused by POMC mutations in humans. Nat Genet. 1998 Jun;19(2):155–7.

33. Yaswen L, Diehl N, Brennan MB, Hochgeschwender U. Obesity in the mouse model of pro-opiomelanocortin deficiency responds to peripheral melanocortin. Nat Med. 1999 Sep;5(9):1066–70.

34. Steculorum SM, Ruud J, Karakasilioti I, Backes H, Ruud LE, Timper K, et al. AgRP Neurons Control Systemic Insulin Sensitivity via Myostatin Expression in Brown Adipose Tissue. Cell. 2016 Mar 24;165(1):125–38.

35. Üner A, Gonçalves GHM, Li W, Porceban M, Caron N, Schönke M, et al. The role of GluN2A and GluN2B NMDA receptor subunits in AgRP and POMC neurons on body weight and glucose homeostasis. Mol Metab. 2015 Oct;4(10):678–91.

36. Suyama S, Yada T. New insight into GABAergic neurons in the hypothalamic feeding regulation. J Physiol Sci. 2018 Nov 1;68(6):717–22.

37. Rozov A, Burnashev N. Fast interaction between AMPA and NMDA receptors by intracellular calcium. Cell Calcium. 2016;60(6):407–14.

38. Ibrahim N, Bosch MA, Smart JL, Qiu J, Rubinstein M, Rønnekleiv OK, et al. Hypothalamic proopiomelanocortin neurons are glucose responsive and express K(ATP) channels. Endocrinology. 2003 Apr;144(4):1331–40.

39. Roberts BL, Zhu M, Zhao H, Dillon C, Appleyard SM. High glucose increases action potential firing of catecholamine neurons in the nucleus of the solitary tract by increasing spontaneous glutamate inputs. Am J Physiol Regul Integr Comp Physiol. 2017 Sep 1;313(3):R229–39.

40. Wu Z-Y, Zhu L-J, Zou N, Bombek LK, Shao C-Y, Wang N, et al. AMPA receptors regulate exocytosis and insulin release in pancreatic β cells. Traffic. 2012 Aug;13(8):1124–39.

41. Hu J, Jiang L, Low MJ, Rui L. Glucose Rapidly Induces Different Forms of Excitatory Synaptic Plasticity in Hypothalamic POMC Neurons. PLoS One [Internet]. 2014 Aug 15 [cited 2019 Apr 2];9(8). Available from: https://www.ncbi.nlm.nih.gov/pmc/articles/PMC4134273/

42. McClelland AC, Hruska M, Coenen AJ, Henkemeyer M, Dalva MB. Trans-synaptic EphB2-ephrin-B3 interaction regulates excitatory synapse density by inhibition of postsynaptic MAPK signaling. Proc Natl Acad Sci USA. 2010 May 11;107(19):8830–5.

43. Cansell C, Denis RGP, Joly-Amado A, Castel J, Luquet S. Arcuate AgRP neurons and the regulation of energy balance. Front Endocrinol (Lausanne) [Internet]. 2012 Dec 27 [cited 2019 Dec 18];3. Available from: https://www.ncbi.nlm.nih.gov/pmc/articles/PMC3530831/

44. Hubbard K, Shome A, Sun B, Pontré B, McGregor A, Mountjoy KG. Chronic High-Fat Diet Exacerbates Sexually Dimorphic Pomctm1/tm1 Mouse Obesity. Endocrinology. 2019 01;160(5):1081–96.

45. Wang C, Xu Y. Mechanisms for Sex Differences in Energy Homeostasis. J Mol Endocrinol. 2019 Feb;62(2):R129–43.

46. Wang C, He Y, Xu P, Yang Y, Saito K, Xia Y, et al. TAp63 contributes to sexual dimorphism in POMC neuron functions and energy homeostasis. Nat Commun. 2018 18;9(1):1544.

47. Lein ES, Hawrylycz MJ, Ao N, Ayres M, Bensinger A, Bernard A, et al. Genome-wide atlas of gene expression in the adult mouse brain. Nature. 2007 Jan 11;445(7124):168–76.

48. He Z, Gao Y, Alhadeff AL, Castorena CM, Huang Y, Lieu L, et al. Cellular and synaptic reorganization of arcuate NPY/AgRP and POMC neurons after exercise. Mol Metab. 2018 Sep 12;

49. Newton AJ, Hess S, Paeger L, Vogt MC, Fleming Lascano J, Nillni EA, et al. AgRP Innervation onto POMC Neurons Increases with Age and Is Accelerated with Chronic High-Fat Feeding in Male Mice. Endocrinology. 2013 Jan;154(1):172–83.

50. Madisen L, Zwingman TA, Sunkin SM, Oh SW, Zariwala HA, Gu H, et al. A robust and high-throughput Cre reporting and characterization system for the whole mouse brain. Nat Neurosci. 2010 Jan;13(1):133–40.

51. van den Pol AN, Yao Y, Fu L-Y, Foo K, Huang H, Coppari R, et al. Neuromedin B and gastrin-releasing peptide excite arcuate nucleus neuropeptide Y neurons in a novel transgenic mouse expressing strong Renilla green fluorescent protein in NPY neurons. J Neurosci. 2009 Apr 8;29(14):4622–39.

52. van der Klaauw AA, Croizier S, Mendes de Oliveira E, Stadler LKJ, Park S, Kong Y, et al. Human Semaphorin 3 Variants Link Melanocortin Circuit Development and Energy Balance. Cell. 2019 07;176(4):729–742.e18.

53. Dobin A, Davis CA, Schlesinger F, Drenkow J, Zaleski C, Jha S, et al. STAR: ultrafast universal RNA-seq aligner. Bioinformatics. 2013 Jan;29(1):15–21.

54. Anders S, Pyl PT, Huber W. HTSeq—a Python framework to work with high-throughput sequencing data. Bioinformatics. 2015 Jan 15;31(2):166–9.

55. Wang L, Wang S, Li W. RSeQC: quality control of RNA-seq experiments. Bioinformatics. 2012 Aug 15;28(16):2184–5.

56. Li, Dewey. RSEM: accurate transcript quantification from RNA-Seq data with or without a reference genome. BMC Bioinformatics volume. 2011;(12):323.

57. Robinson MD, McCarthy DJ, Smyth GK. edgeR: a Bioconductor package for differential expression analysis of digital gene expression data. Bioinformatics. 2010 Jan 1;26(1):139–40.

58. Croizier S, Prevot V, Bouret SG. Leptin Controls Parasympathetic Wiring of the Pancreas During Embryonic Life. Cell Rep. 2016 Apr 5;15(1):36–44.

59. Magnan C, Collins S, Berthault M-F, Kassis N, Vincent M, Gilbert M, et al. Lipid infusion lowers sympathetic nervous activity and leads to increased β-cell responsiveness to glucose. J Clin Invest. 1999 Feb 1;103(3):413–9.

60. Picard A, Soyer J, Berney X, Tarussio D, Quenneville S, Jan M, et al. A Genetic Screen Identifies Hypothalamic Fgf15 as a Regulator of Glucagon Secretion. Cell Rep. 2016 08;17(7):1795–806.

61. Tarussio D, Metref S, Seyer P, Mounien L, Vallois D, Magnan C, et al. Nervous glucose sensing regulates postnatal β cell proliferation and glucose homeostasis. J Clin Invest. 2014 Jan;124(1):413–24.

